# Predicting the direction of phenotypic difference

**DOI:** 10.1101/2024.02.22.581566

**Authors:** David Gokhman, Keith D. Harris, Shai Carmi, Gili Greenbaum

**Author notes:** Equal contribution (David Gokhman, Keith D. Harris).

## Abstract

Predicting phenotypes from genomic data is a key goal in genetics, but for most complex phenotypes, predictions are hampered by incomplete genotype-to-phenotype mapping. Here, we describe a more attainable approach than quantitative predictions, which is aimed at qualitatively predicting phenotypic differences. Despite incomplete genotype-to-phenotype mapping, we show that it is relatively easy to determine which of two individuals has a greater phenotypic value. This question is central in many scenarios, e.g., comparing disease risk between individuals, the yield of crop strains, or the anatomy of extinct vs extant species. To evaluate prediction accuracy, i.e., the probability that the individual with the greater predicted phenotype indeed has a greater phenotypic value, we developed an estimator of the ratio between known and unknown effects on the phenotype. We evaluated prediction accuracy using human data from tens of thousands of individuals from either the same family or the same population, as well as data from different species. We found that, in many cases, even when only a small fraction of the loci affecting a phenotype is known, the individual with the greater phenotypic value can be identified with over 90% accuracy. Our approach also circumvents some of the limitations in transferring genetic association results across populations. Overall, we introduce an approach that enables accurate predictions of key information on phenotypes — the direction of phenotypic difference — and suggest that more phenotypic information can be extracted from genomic data than previously appreciated.

## Introduction

A key goal in genetics is to predict phenotypes from genomic data. Such predictions are pivotal for assessing disease risk (1; 2), understanding the genetics underlying adaptation (3; 4; 1), optimizing genetic engineering outcomes (5), reconstructing the traits of extinct species (6), and more. However, our current ability to predict phenotypic values from genetic information, for example by using polygenic scores (PGSs), is restricted by several factors. These include environmental effects, the high polygenicity of many phenotypes, the limited ability to identify causal noncoding variants and quantify their effects, and the lack of power to detect small-effect loci (1; 2).

Given the limitations associated with predicting precise phenotypes, we suggest here a more attainable objective: predicting only the direction of phenotypic difference. Namely, rather than striving to predict the precise phenotypic value encoded by a particular genome, we aim to predict whether this genome encodes for a larger or smaller phenotypic value relative to another genome. To illustrate, consider a scenario where one is interested in determining the probability that an offspring will be taller than their 170cm tall parent. Considering that a PGS predicts the offspring will be 180cm tall, what is the probability that the offspring will indeed be taller than their parent? We previously implemented a simplified version of this approach to reconstruct Denisovan anatomy using gene regulatory data, and validated the method on Neanderthals and chimpanzees, finding that it reaches over 85% accuracy in predicting the direction of phenotypic differences (6).

Undoubtedly, predicting a precise phenotypic value is more informative than predicting only the direction of phenotypic difference. However, in studies where the precise phenotypic value cannot be accurately inferred (which is often the case), important insights could still be gained by inferring the phenotypic direction instead. Most importantly, the phenotypic direction is often the crux of phenotypic comparisons, for example, when estimating how likely it is that (i) an individual has an increased disease risk compared to a reference (2), (ii) a genetically modified crop will have increased yield (7), (iii) an individual will be greater or smaller in a certain trait compared to their parents or siblings (e.g., in preimplantation genetic diagnoses; (8; 9)), and (iv) the phenotypes of an extinct species differ from those of an extant species.

Here, we explored the feasibility of using currently available genotype-to-phenotype information to predict which individual has a greater phenotypic value. We compared the total effect of known loci to the range of the potential effects of unknown genetic and non-genetic contributors. We studied this ratio of known-to-unknown effects through two independent branches of investigation: (i) formalizing a model to delineate the scenarios in which accurate predictions can be achieved, and (ii) evaluating performance in real-world empirical data from humans and other species, examining a wide range of levels of divergence between individuals. Our findings underscore the known-to-unknown ratio as a high-fidelity and intuitive estimator of prediction accuracy. This allowed us to identify cases where we can reliably discern the individual with the greater phenotypic value. Importantly, this is possible even in cases where the proportion of variance in the trait explained by known genetic effects is small. Our study suggests that it is possible to identify the pairs of individuals for which high-accuracy predictions can be made, and that more phenotypic information can be reliably extracted from a genome than perhaps intuitively expected.

## Results

### Approach

We investigated what genomic information is needed to predict the direction of phenotypic difference between two individuals and the conditions under which this prediction is accurate. We assume that one individual has been phenotyped (hereafter, *phenotyped* individual) and the other has not (hereafter, *unphenotyped* individual).

A phenotype is affected by loci whose contribution to (or often association with) the phenotype is known (hereafter, *known effects*), as well as by loci or non-genetic factors whose association with the phenotype is unknown (hereafter, *unknown effects*, Fig. 1a). We make a prediction on the direction of phenotypic difference by summing up the contribution of the known effects and determining whether the unphenotyped individual has a larger or a smaller sum. We ignore loci where the two compared individuals have the same genotype, because only divergent loci could contribute to the phenotypic difference (Fig. 1b,c). This procedure is equivalent to computing the difference between the PGSs of the two genomes, and using the sign of this difference to predict the direction of the phenotypic difference (9; 10). In the following sections, we investigated the conditions affecting the probability that a prediction based only on the known effects matches the true direction of phenotypic difference (hereafter, *prediction accuracy* or *P*).

**Figure 1:**
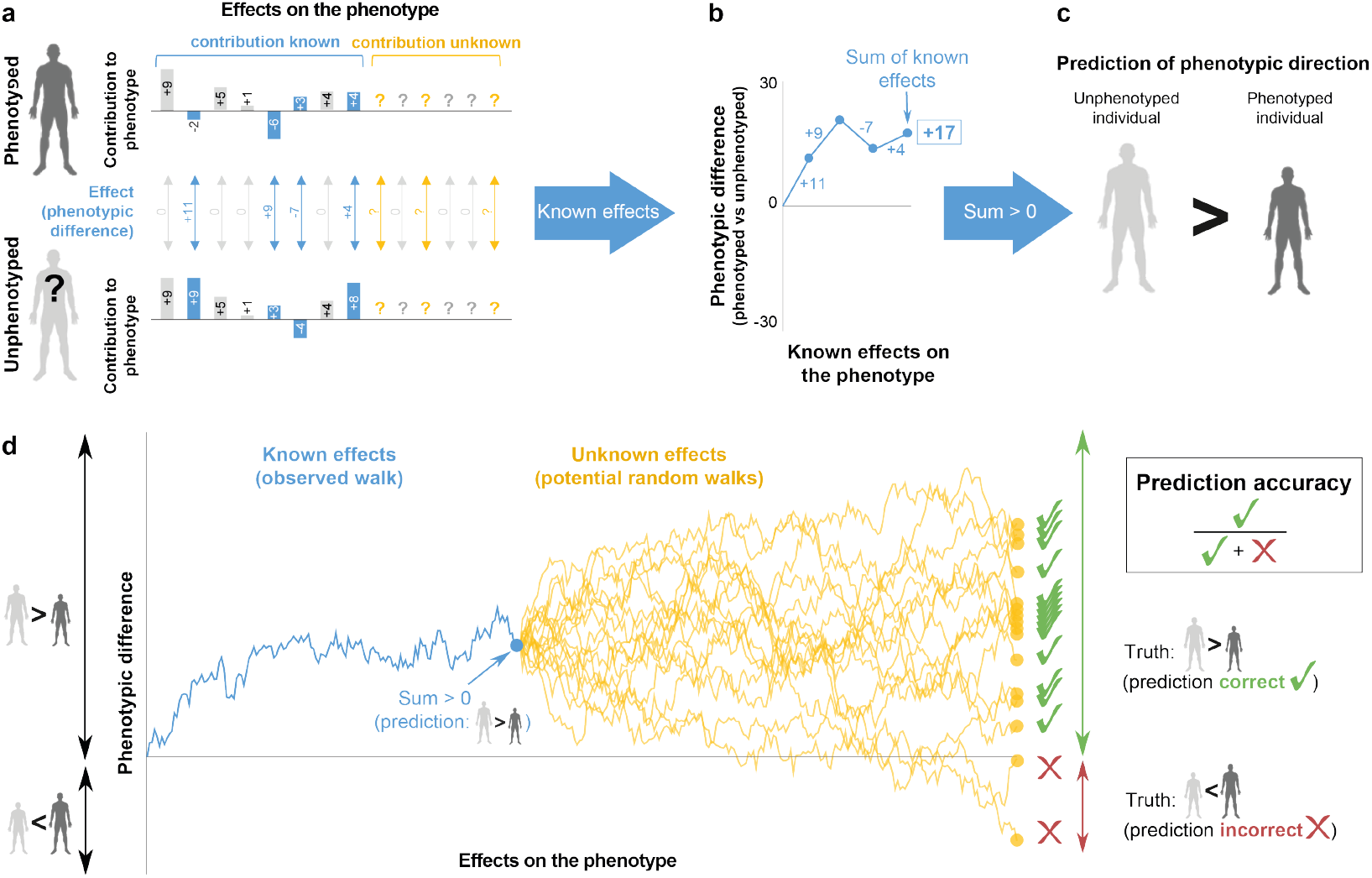
Schematic of the approach to predict the direction of phenotypic difference. (a) We start with a phenotyped individual and an unphenotyped individual. We consider the known and unknown effects contributing to (or associated with) the phenotype of interest. Known genetic effects on the phenotypic difference are in blue (measured in units of the phenotype), unknown genetic and non-genetic effects are in yellow. Cases where the contribution is identical between the two individuals (and therefore do not affect the phenotypic difference) are in gray. (b) Only the known divergent effects are used to predict the phenotypic difference between the individuals. The sum of the known effects can be thought of as the final position of a random walk with step sizes and directions corresponding to the effect sizes. (c) The direction of the total sum of the known effects is used to make a prediction of the direction of phenotypic difference between the phenotyped and unphenotyped individuals. If the sum of the known effects between the individuals is positive, we predict that the phenotypic value of the unphenotyped individual is larger than the phenotyped individual (and the opposite prediction if the sum is negative). (d) Modeling prediction accuracy using random walks. The curves represent random walks where each step is an effect size. The blue curve shows the known effects of a specific random walk, and the sign (positive or negative) of the blue point at the end of the walk is the predicted direction of phenotypic difference. The yellow curves show potential random walks of the unknown effects (genetic and environmental). In this example, effect sizes were drawn from a standard normal distribution. For a correct prediction of the direction of the phenotypic difference, the sum of the known effects (blue point) and the true phenotypic difference (yellow dot) need to be on the same side of the x-axis (both below or both above).

### Modeling the conditions needed to predict the phenotypic direction

We explored the problem from two different perspectives, statistical genetics and evolutionary, which provide different tools and intuitions. From a statistical genetics perspective, we considered the partitioning of the phenotypic variance into that generated by known and unknown effects. For the evolutionary perspective, we modeled the approach as a random walk, where each step is an effect on the phenotype in one or the other direction. We define the effect size of a locus as the average difference in predicted phenotype between the genotypes of the two individuals. For example, if the phenotyped individual has a genotype that increases height by 3mm (relative to a reference), and the unphenotyped individual has another genotype, which decreases height by 1mm, then we consider the effect size of that locus to be +4mm (Fig. 1a). The effect size of loci with the same genotype in the two individuals is 0, and these loci are therefore ignored throughout this work. Our model makes the simplifying assumptions of additivity and no epistasis (11) (in the empirical section, where we test our approach, these simplifying assumptions are evaluated). The direction of the sum of known effects (i.e., whether the displacement is above or below the x-axis in Figure 1b and the blue dot in Figure 1d) is our prediction of the direction of the phenotypic difference (Fig. 1c). If the remaining steps of the random walk (i.e., those of the unknown effects) are such that the final displacement (i.e., true phenotype, yellow dots in Fig. 1d) is still above 0, our prediction is correct. Otherwise, i.e., if the remaining steps push the displacement below 0, our prediction based on the known effects is incorrect. Naturally, the larger the sum of known effects is, the less likely it is for the final displacement to end on the opposite side of the x-axis.

We start by exploring the factors affecting prediction accuracy and the conditions required for high-accuracy predictions. Various factors have the potential to affect prediction accuracy: the total number of loci affecting a phenotype, the fraction of known effects, the distribution of effect sizes, and more. However, our random walk perspective suggests that all of these factors amount to only two aspects of the walk that ultimately determine prediction accuracy. The first aspect is the vertical displacement of the sum of the known effects (blue dot in Fig. 1d; equivalent in statistical genetics to the difference in PGS). Namely, the further above or below 0 we “traveled”, the less likely it is that the unknown effects would push the final position to the other side of the x-axis. The second aspect is the variation of the overall potential sums of the unknown effects (i.e., the variation in the displacements generated by the random walk of the unknown effects, yellow region in Fig. 1d; equivalent to the proportion of variance in phenotypic differences that is unexplained by PGS). The smaller this variation is, the less likely the unknown effects are to push the final position of the walk to the other side. We propose here that prediction accuracy can be characterized by the ratio between these two quantities. Denoting the sum of the known effects as Δ and the standard deviation of the unknown effects as *σ*, we define the *known-to-unknown ratio, κ*, as

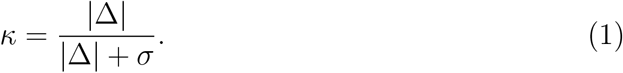

In *Methods*, we show that the prediction accuracy can be written as a simple function of *κ*,

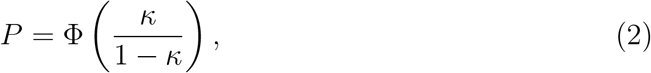

where Φ(*·*) is the standard normal CDF. We provide two derivations — one from the viewpoint of random walks and the other from the viewpoint of statistical genetics, which also enabled us to model shared genetic and environmental components in siblings (see *Methods*). We then explored how different factors affect the distribution of *κ*, by deriving the distributions under simplified conditions (*Supplementary Information*) as well as using simulations. We simulated pairs of individuals with random known and unknown effects and arbitrarily treated one individual as phenotyped and the other as unphenotyped (see *Methods*). Based on these simulated effects, we computed *κ* for each pair of individuals and determined whether the prediction is correct. We conducted these comparisons for different ratios of known to unknown effects, as well as for different effect-size distributions.

We found an agreement between the theoretical expectation and the simulated results across all values of *κ* (Fig. 2a), as well as across different effect-size distributions (Fig. S1a–b). As expected, predictions on pairs with higher *κ* values showed higher prediction accuracy. For example, for pairs of individuals with *κ >* 0.62, prediction accuracy was *P >* 0.95. High values of *κ* are more common when the fractions of known effects are larger (Fig. 2b), but we showed analytically (*Supporting information*) and with simulations (Fig. S1c–d) that the underlying effect-size distribution does not affect the *κ* distributions (Fig. S1c–d).

**Figure 2:**
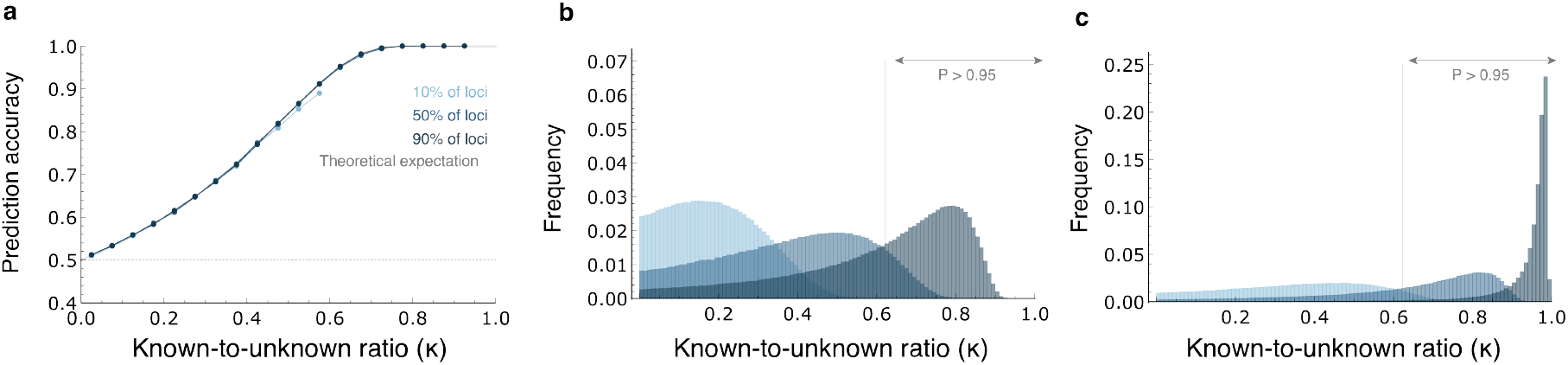
Evaluating prediction accuracy using the known-to-unknown ratio (*κ*). (a) Simulated prediction accuracies for various *κ* values (grouped into equally spaced bins), for different proportions of the known vs. unknown effects (10%, 50%, and 90% of effects known). Effect sizes were drawn from a normal distribution. In gray is the theoretical expectation from Eq. 4. (b) The distribution of *κ* values for the case where the known effects are randomly sampled. The vertical line denotes the *κ* values required for prediction accuracy of *P >* 0.95 (*κ* = 0.62) (c). The distribution of *κ* values for the case where the known effects are those with the largest effect sizes. The vertical line denote the *κ* values required for prediction accuracy of *P >* 0.95. In all panels, 10,000 effect sizes were drawn from a standard normal distribution to represent the known and unknown effects on the phenotype.

We have so far assumed that there is no bias in choosing which effects are known and which effects are unknown. However, many detection methods (e.g., quantitative trait loci mapping or GWAS) have an ascertainment bias, where loci with larger effects are more readily detectable (2). We therefore analyzed scenarios where the known effects are those with the largest contribution to phenotypic variance. As before, we found that *κ* is a precise descriptor of prediction accuracy (Fig. S2). However, *κ* values tend to be much higher than in the unbiased scenario (Fig. 2c). Therefore, if the known effects tend to be the largest effects, prediction accuracy could be high. For example, with 10% of effects known in the unbiased scenario, none of the simulated pairs of individuals had prediction accuracy *>* 0.95 (*κ >* 0.62); however, in the scenario where the largest effects were known, 6.5% of the pairs reached this prediction accuracy (Fig. 2b–c, intermediate blue). Thus, if the known effects tend to have larger effects, high prediction accuracy can be achieved even in cases where these loci explain only a small proportion of the overall phenotypic variance.

In sum, we found that the known-to-unknown ratio (*κ*) captures the factors that affect the probability of predicting which individual has the higher phenotypic value. The *κ* estimator could thus be used as an intuitive statistic to (i) evaluate prediction accuracy, and (ii) identify individuals for which high-accuracy predictions could be made, even when genotype-to-phenotype data is limited.

### Identifying which individual has the higher phenotypic value in real-world data

To investigate the relationship between *κ* and prediction accuracy in empirical data, we compared pairs of individuals with different levels of genetic divergence. We considered pairs of individuals from the UK Biobank (12) from either the same family or the same population. For each pair, we investigated six phenotypes: height, body mass index (BMI), metabolic rate, blood pressure, hip circumference, and bone density. For each phenotype, we selected loci that significantly contribute to the phenotype based on a GWAS that excluded the individuals we tested. The effect sizes generated in this GWAS were then used to compute Δ as the difference between the PGSs of the two individuals (*Methods*). In each comparison, we also computed *κ*. For the within-family comparisons, we examined all 10,597 pairs of same-sex siblings in the dataset (*Methods*). For within-population comparisons, we randomly sampled 20,000 individuals (10,000 females and 10,000 males) who self-identified as White British and had Northwestern European genetic ancestry (hereafter labeled for brevity as ‘European’, see Methods, Fig. S6). We then examined all pairwise same-sex comparisons among them.

Across the six phenotypes, higher *κ* values reflected higher prediction accuracy (Fig. 3a–b), with a relationship that tightly followed the theoretical expectation (Eq. 4). Importantly, this is maintained across both levels of genetic divergence between individuals (family-level and population-level), suggesting that *κ* captures the key aspects determining the ability to predict phenotypes. There is an intriguing exceptions to this: predictions of blood pressure differences hold at lower *κ* values, but perform badly at higher *κ* values. This possibly reflects medication-induced phenotypic changes (see below).

**Figure 3:**
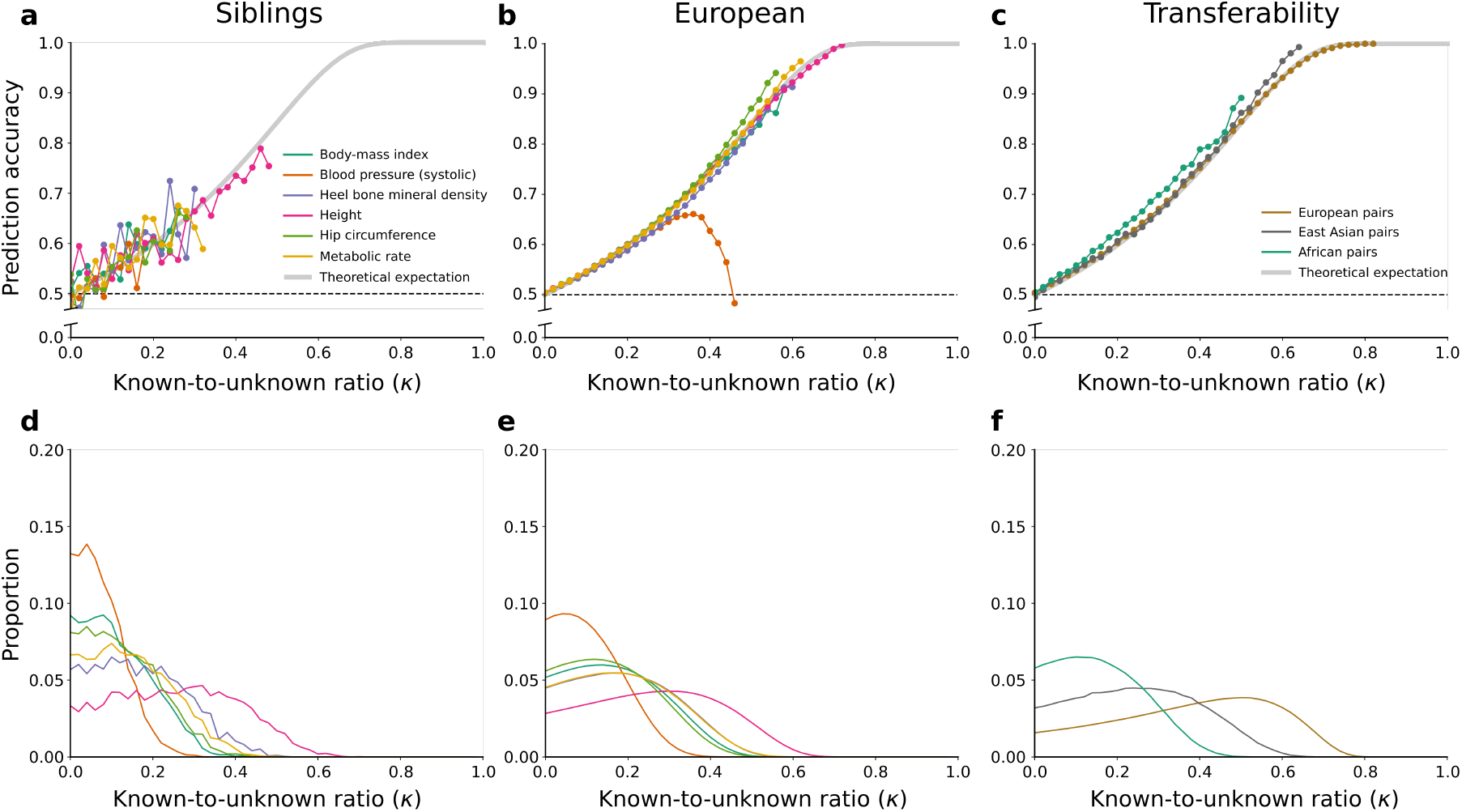
Predictions of the direction of phenotypic difference in humans. (a)–(c) The relationship between the known-to-unknown ratio (*κ*) and the proportion of correct predictions in different phenotypes. The theoretical expectation (Eq. 4) is shown in gray. (a) Pairwise comparisons of siblings from the UK Biobank for six phenotypes. (b) Pairwise comparisons of individuals from the European group (self-identified White British with Northwestern European genetic ancestry) from the UK Biobank for the same six phenotypes. (c) Pairwise height comparisons of individuals from the same population (either European, East Asian or African, as defined in Fig. S6), using GWAS generated from a European-ancestry group in Yengo *et al*. (15). (d)–(f) The distribution of *κ* values for all pairwise comparisons. Each panel corresponds to the panel above it.

Our approach also allowed us to estimate the proportion of individuals for whom high-accuracy predictions can be achieved. For example, for 5% of pairs from the European group, *κ* values for bone mineral density are *≥* 0.4, and we can therefore predict which individual has higher bone mineral density with 75% accuracy (i.e., threefold more likely to predict correctly than incorrectly; Figs. 3e and S4b). For height, where a larger fraction of loci contributing to the phenotypic variance is known, the same prediction accuracy can be achieved for one in four pairs. Notably, we can predict the taller individual with 90% certainty for 3% of the pairs (Fig. S4a). Importantly, the percentage of pairs for which high-accuracy predictions can be attained increases with increasing genetic distance (*κ* distributions are shifted to the right with higher divergence between pairs in Fig. 3d–e). For example, in 3% of sibling pairs, we can predict which sibling is taller with 85% certainty, and between unrelated individuals from the European group, this increases to 8% of pairs (Fig. S4a–b). It remains to be determined to what extent these results are affected by population stratification (13) or other potential factors.

One of the most intriguing uses of phenotypic inference is its potential to predict an individual’s susceptibility to a particular disease. Since disease risk is not directly quantifiable per individual, we tested instead our ability to identify the individual with the disease in a pair of individuals where one is healthy and the other is reported to have the disease. Here too, the empirical results mostly align with the theoretical expectation. However, unlike all other analyses, at higher *κ* values (*κ >∼* 0.4), the empirical results started to deviate from the theoretical expectation (Fig. S5a). We have not been able to pinpoint the underlying driver of this phenomenon. One plausible explanation is that in these comparisons, higher *κ* values reflect instances where one of the individuals is indeed more likely to develop the disease, but early signs of the disease or family history prompted treatment and thus exclusion from the disease group. Potential support for this can be seen in the context of the blood pressure phenotype. At higher *κ* values, predictions start diverging from the theoretical expectation both in the within-population analysis of blood pressure (Fig. 3b), as well as in the disease analysis of hypertension (Fig. S5a), where for high *κ* values prediction accuracy approaches 0 and thus our predictions are not even random, but systematically wrong. This behavior may indicate a negative correlation between high *κ* values and the disease, possibly reflecting medication-induced phenotypic changes that specifically affect individuals with a higher likelihood of elevated blood pressure, thereby altering the predictive outcome. Nevertheless, for most cases, where *κ* values are not extreme, it is possible to generate accurate estimates of prediction accuracy. This could perhaps be clinically relevant when the unphenotyped individual has a higher probability of developing the disease relative to an individual known to have the disease.

A major concern in GWAS is its limited transferability across populations. PGSs computed using data from one population often perform substantially worse when applied to other populations (14). To test whether this phenomenon affects our approach, we evaluated the relationship between *κ* and prediction accuracy using GWAS conducted on individuals with European ancestry, but predicting phenotypes between pairs of individuals with East Asian or African Ancestry (populations defined in (15)). As expected, we observed lower *κ* values for these comparisons relative to the *κ* distribution in Europeans (Fig. 3f), highlighting that prediction accuracy in non-European populations is worse than in Europeans, owing to the smaller fraction of the phenotypic variance explained by European-ancestry GWASs (14; 15). This, in turn, may lead to inequality in future gains from genomics-based medicine. Nevertheless, here too, we observed good agreement with the theoretical expectation for the relationship between *κ* and prediction accuracy (Fig. 3c). Thus, while fewer usable SNPs and increased noise in effect size estimation lead to fewer pairs with high-accuracy predictions, the ability to robustly estimate prediction accuracy is maintained.

In summary, we found that: (i) given a pair of individuals, we are able to accurately estimate the chances of correctly predicting which individual has the greater phenotypic value, and (ii) even for phenotypes with limited genotype-to-phenotype data, some pairs have sufficiently high known-to-unknown ratios (*κ*) to enable the identification of the individual with the greater phenotypic value. Two important implications of these findings are that we can (i) select the subset of pairs of individuals for which we can make high-confidence predictions, or (ii) given a pair of individuals, select the subset of phenotypes for which we can make high-confidence predictions.

### Impact of directional selection on predictions between populations and species

In the model above, we have not addressed the role of selection. Directional selection most likely has little effect on the within-population UK Biobank comparisons, but may play a more central role when more divergent genomes are compared. In this section, we extend our model to include directional selection and examine predictions in divergent populations and species (see *Discussion* for the potential effects of negative and stabilizing selection).

Until now, our model assumed that the effects have an equal probability of increasing or decreasing the phenotypic difference. Under directional selection, the phenotype of a lineage is typically pushed towards a new optimal value. The directions of effects of that lineage relative to the ancestral lineage are more likely to be in the direction of this optimum (16). Thus, to model the case that directional selection has shaped the divergence between the two compared genomes, we introduced biased effects into our model. We considered the case where selection is stronger for larger effect sizes. In other words, effects are more likely to be aligned with the direction of selection than with the opposite direction, and the probability of alignment increases with the size of the effect and the strength of selection.

To model this, we introduced into the random walk a bias that favors one direction over the other and is stronger with larger effects (*Methods*). In this model, we observed an improvement in prediction accuracy relative to the neutral case in two aspects: (i) the proportion of pairs of individuals with high *κ* values also increases with stronger selection (Fig. S3); (ii) prediction accuracy is higher for any given value of *κ* (Fig. 4b). Both improvements increase with stronger directional selection. Consequently, under directional selection, high-accuracy predictions can be achieved more often and with fewer known effects.

**Figure 4:**
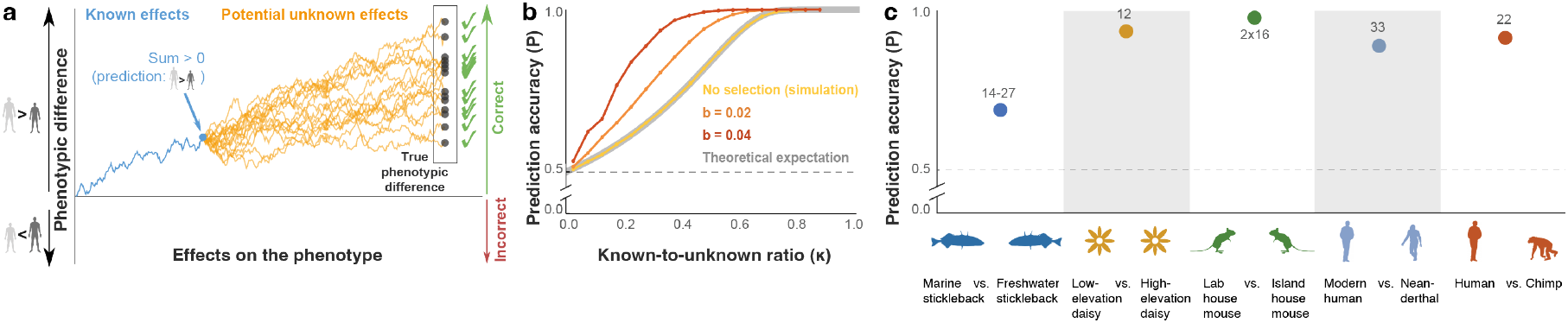
The effect of directional selection on predicting the direction of phenotypic difference. (a) Prediction accuracy under directional selection, modeled as a biased random walk. The random walks in this schematic are biased toward the positive direction, with larger effects having a stronger bias. Biased random walks increase prediction accuracy. (b) Prediction accuracy for different *κ* values and different levels of bias, with 50% randomly selected known effects out of 10,000 overall. (c) Prediction accuracy across species. Each point represents the proportion of correct predictions. The number of phenotypes is noted above each data point. For sticklebacks, between 14 and 27 phenotypic predictions were made for four different freshwater populations. For mice, predictions were made for two phenotypes in 16 developmental stages.

These results suggest that more divergent lineages, where directional selection might have played a more central role, would tend to show higher prediction accuracy. To investigate this, we explored genotype-to-phenotype datasets of more divergent lineages. We tested three quantitative trait loci (QTL) mapping datasets investigating stickleback (17), daisy (18), and mouse (19). The stickleback dataset included four freshwater populations that diverged from a common marine ancestor less than 12,000 years ago (17). We analyzed the 27 morphological phenotypes in the dataset, with 1–2 QTLs reported per phenotype, and found that even with only 1-2 known loci, prediction accuracy was 63%-75% (depending on the pair of populations compared; Fig. 4c).

In the daisy dataset, we analyzed 1–5 QTLs for 12 phenotypes that differ between two species of daisy (18). We found a prediction accuracy of 92%, with 11 out of 12 phenotypes predicted correctly based on these known effects (Fig. 4c). The mouse dataset included growth rate and weight phenotypes of Gough Island vs. wild-type mice over 16 developmental stages (19), with 8–11 QTL per phenotype. Prediction accuracy was 100% (Fig. 4c). Interestingly, this perfect prediction accuracy is achieved despite the fact that in some developmental stages, the joint effect of all known effects explains as little as 6% of the variance in weight and 3% of the variance in growth rate. In addition, in all three datasets, the single largest-effect locus was sufficient to predict the direction of phenotypic difference with high accuracy (63%-75% for sticklebacks, 92% for daisies, and 75% for mice).

We also revisited our previous study that predicted phenotypic differences between Neanderthals and modern humans and between chimpanzees and humans (6). These predictions were based on DNA methylation data and were made only for phenotypes where all known effects pointed in the same direction of phenotypic change, thus filtering for phenotypes with higher *κ* values. Prediction accuracy for 33 Neanderthal phenotypes and 22 chimpanzee phenotypes was 88% and 91%, respectively (6). Interestingly, we observed similar patterns in our more recent study comparing human and chimpanzee gene expression in human-chimpanzee hybrid cells, with an accuracy of 81% (20).

Overall, these datasets represent a diverse range of phenotypes, species, divergence times, and genotype-to-phenotype association methods. While we most often do not know the exact nature of the selection processes that have shaped the genetics of organisms, our results suggest that when comparing divergent genomes, we can achieve relatively accurate prediction of the direction of the phenotypic difference with very few large-effect loci.

## Discussion

Traditional quantitative genetic studies attempt to predict the precise phenotypic value of an individual. Here, we explored a more modest approach, whereby only the direction of phenotypic difference is predicted. Our goal was to develop a model for prediction accuracy under various conditions and to test it on empirical data. We found that prediction accuracy is affected by two main factors: the sum of known effects, and the variance of the sums of the unknown effects. We formulated the relationship between these two factors as *κ*, from which the prediction accuracy can be easily estimated. The *κ* statistic allows us to identify pairs of individuals where the direction of phenotypic difference could be confidently predicted. This statistic is not affected by ascertainment bias, the level of divergence between individuals, or transferability problems with the data. Pairs for whom accurate predictions can be made are more common when (i) more information is known about the genetic basis of the phenotypic variation, (ii) the phenotype was more strongly affected by positive selection, (iii) large-effect loci are more likely to be known.

Our model has several limitations. (i) We assumed additivity of effect sizes and did not incorporate epistasis. Although previous studies have shown that variation in complex traits within species is mostly additive (21; 22; 23), the assumption of additivity may not hold for some phenotypes (24; 25). (ii) In our model, we did not separate between unknown effects that contribute to the phenotype (e.g., undetected loci) and unknown effects due to noise in the estimation of known effects (e.g., measurement errors or unaccounted factors such as age and socio-economic status). (iii) Finally, we model environmental effects as part of the unknown effects, i.e., reflecting the same dynamics. However, in phenotypes that evolve under stabilizing selection in the face of shifting environments, genetic and environmental effects can have different or opposing trends (26; 27). Despite these limitations, testing our approach on real data suggests that our current model captures the main factors affecting predictions. To model selection, we used an approach where loci are affected by selection in proportion to their effect sizes. While this is the general case, selection often follows more complex dynamics (3). For example, Hayward and Sella (16) investigated temporal evolutionary dynamics of a rapid adaptation phase followed by a prolonged stabilizing selection phase. This study showed that in the long term, phenotypic variation is dominated primarily by small and moderate effect sizes, and that the larger the effect size of a locus that separates the two groups, the more likely it is to reflect the overall phenotypic difference between them (16). This could further explain the high prediction accuracy reached in our between-species comparisons, where the few known large-effect loci explain a small percentage of the overall phenotypic difference, but are very predictive of the direction of phenotypic difference.

Other types of selection could also affect predictions. For example, negative selection is expected to reduce the number of divergent loci between two individuals, thus decreasing both the known and unknown effects. If it disproportionately affects larger-effect loci it might reduce the relative contribution of the known effects, thus shifting *κ* values towards lower values, resulting in lower prediction accuracy. Unlike directional selection, this is not expected to affect the relation between *κ* and prediction accuracy. Stabilizing selection, for a similar optimum on the two genomes, may also reduce prediction accuracy because it can reduce the variance contribution of shared loci affecting the phenotype (27).

The ability to predict the direction of some phenotype differences accord with recent results in the context of embryo screening for polygenic conditions. In this setting, embryos generated by *in vitro* fertilization are screened for their genetic predisposition to complex, polygenic diseases and ranked for transfer for pregnancy. This technology is considered by many as unethical (28; 29; 30) and is also widely believed to have little ability to identify embryos with substantially lower risk (31; 32). However, several simulation and modeling studies (9; 10; 33; 34) found that even for diseases with poorly predictive PGS, selection of the lowest-risk embryo could lead to substantial relative risk reduction. This result can be understood based on the liability threshold model (35), under which a disease is assumed to have an underlying, unobserved, continuous liability, with affected individuals being those with liability exceeding a threshold. Given a weak PGS, embryo screening would reduce the expected liability of the selected embryo by a very small amount compared to a randomly selected embryo (8; 36). However, even this small shift could be sufficient to substantially reduce the chances of exceeding the disease threshold. Similarly, when predicting the direction of the phenotypic difference, even a small positive gap in PGS between a pair of individuals translates to a high probability for the final phenotypic difference to remain positive.

The approach we presented evaluates the extent to which a key feature of a phenotype — its direction — can be predicted from genomic data. Given the currently limited ability to quantitatively predict phenotypes from genotypes (2), our approach suggests that qualitative prediction of phenotype direction is often feasible. While there is still much to explore with regard to the applicability of this approach to various data, its capability to robustly estimate prediction accuracy and to identify individuals and phenotypes for which accurate predictions can be achieved, suggests that more phenotypic information can be extracted from genomes than previously appreciated.

## Methods

### Formal model for prediction accuracy

We consider a pair of individuals, one phenotyped and the other unphenotyped, with genomes that diverge at *n* loci that affect a certain phenotype. We denote the (absolute value of the) differential effect of these loci as *e*_*i*_, (*i* = 1, …, *n*) which is the relative contribution of locus *i* to the difference between the phenotypes of the two individuals (Fig. 1a). Each effect of a divergent locus either increases the phenotypic difference in the direction of the phenotyped individual, arbitrarily denoted as *d*_*i*_ = 1, or in the direction of the unphenotyped individual, denoted as *d*_*i*_ = *−*1. The sum of the known effects is 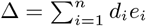 (Fig. 1b). The sign of Δ is our prediction for the direction of the phenotypic difference (Fig. 1c).

We consider additional *m* unknown effects on the phenotype, and denote them as random variables *X*_1_, …, *X*_*m*_. For the most part of this work (but see simulations with selection below), we assume that *X*_1_, …, *X*_*m*_ are independent random variables that attain one of two values, *E*_*j*_ or, *−E*_*j*_, with equal probability, i.e. 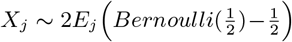, for *j* = 1, …, *m*. We assume that the *E*_*j*_ ‘s are identical independent random variables with an effect-size distribution *Y*, which means that *X*_1_, …, *X*_*m*_ are also identical and independent. Each divergent unknown effect has some contribution to the phenotype, and it can work to either increase or decrease the phenotypic difference. We denote the sum of the unknown effects as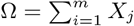.

Following the definitions in Eq. 1, we denote the variance of Ω as *σ*^2^.

The true phenotypic difference is *D* = Δ + Ω, the sum of both known and unknown effects. Our prediction is correct if the signs of Δ and *D* are the same; otherwise, our prediction is incorrect. We define the ‘prediction accuracy’ *P* as the probability that the signs of Δ and *D* are the same.

### Mathematical relationship between *κ* and *P*

Without loss of generality, let us assume that Δ *>* 0. Prediction accuracy is the probability that the true phenotypic difference is positive, *P* = *Prob* (Δ + Ω *>* 0). Reformulating this by plugging in Eq. 1 to replace Δ, we have

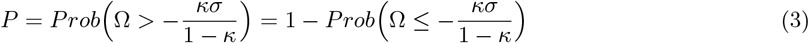

Notably, Ω is a sum of identical independent random variables, and therefore, assuming that the effect size distribution *Y* has a finite variance, we can apply the central limit theorem and show that Ω is approximately normally distributed. Ω has a mean of zero because each of the random variables *X*_*i*_ has a zero mean. We can now use the CDF of Ω, 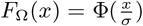(where Φ(*·*) is the standard normal CDF) to explicitly compute the prediction accuracy,

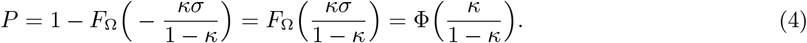

### Alternative derivation

We can also derive this result using standard notations in statistical genetics. As before, we consider that a phenotype is measured in normalized units, i.e., *y ∼ N* (0, 1). The PGS of an individual *p* is then distributed as *p ∼ N* (0, *r*_2_), where *r*_2_ is the proportion of the phenotypic variance explained by the PGS. We denote the combined non-measured genetic factors and non-genetic factors affecting the trait as *e*, which is also the residual of the regression of the trait on the PGS. We can thus write *y* = *p* + *e*. We assume *p* and *e* are independent and *e ∼ N* (0, 1 *− r*^2^). Next, we consider two unrelated individuals with computed PGSs *p*_1_ and *p*_2_ such that *p*_1_ *> p*_2_, with residuals *e*_1_ and *e*_2_, respectively (we assume that *e*_1_ and *e*_2_ are independent because the individuals are unrelated). Denoting the difference in PGSs as *d* = *p*_1_ *− p*_2_, and using *d* to predict the direction of phenotypic difference, the prediction accuracy is therefore *P* = *Prob*(*y*_1_ *> y*_2_), where *y*_1_ and *y*_2_ are the true phenotypic values of the two individuals. We can reformulate this probability as *P* = *Prob*(*e*_2_ *− e*_1_ *< p*_1_ *− p*_2_), and therefore *P* = *Prob*(*e*_2_ *− e*_1_ *< d*). We denote *e*^*′*^ = *e*_2_ *− e*_1_, and because *e*_1_ and *e*_2_ are each normally distributed with variance 1 *− r*^2^ and zero mean, we have *e*^*′*^ *∼ N* (0, 2(1 *− r*^2^)).

We can now observe that:

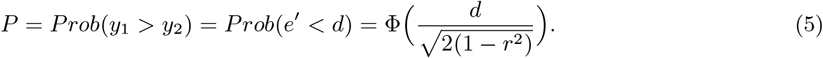

Reformulating Eq. 1 with the notation of this section (i.e., |Δ| = *d* and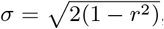, because 2(1 *− r*^2^) is the variance of the differences of the unknown effects), we have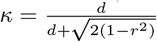, and therefore

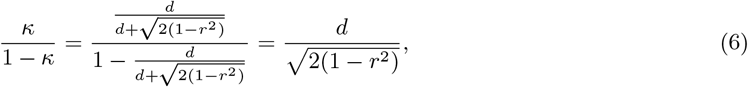

showing that equations 4 and 5 are equivalent.

In the Supporting Information we discuss similar derivations for two specific cases: comparison of siblings and comparison of disease phenotypes.

### Simulations

To simulate a single pairwise comparison, we sampled *n* + *m* effect sizes from a pre-specified effect size distribution, with signs simulated to be negative or positive with equal probability. We then computed the sums 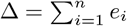 and 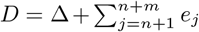, as in the formulation above. The simulation results in a correct prediction if Sign[*D*] = Sign[Δ], otherwise the prediction is incorrect. For each scenario 10^6^ repeats were simulated.

We evaluated different fractions of known effects out of all effects: 10%, 50%, and 90%. Effect size distributions can be shaped by various evolutionary processes, such as mutation, selection, and genetic drift (3; 37); therefore, we simulated effect size distributions of various types (normal distribution in Fig. 2, gamma and Orr’s negative exponential model distributions (38; 3) in Fig. S1). We also considered the case where the known effects tend to be the larger effects. To simulate this, we sampled *n* + *m* effect sizes from the predefined effect size distribution, and then sorted the effect sizes in decreasing order, defining the known effects to be the largest *n* effects. We then continue with the rest of the simulation as described above.

### Modeling and simulating directional selection

To model directional selection, we modify the random variables representing the effects to have positive means. We implement this by simulating *n* + *m* effect sizes *e*_*i*_ as before, but we simulate their direction by letting the probability *X*_*i*_ *>* 0 be 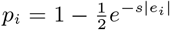, and then *X*_*i*_ *∼* 2*e*_*i* 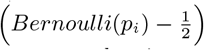_. Note that *s* is not a selection coefficient in units of fitness, but is rather a unitless parameter that is proportional to the impact of selection on the direction of the effect. The motivation for this particular formulation is based on the Ornstein-Uhlenbeck model, which is used to model the evolution of quantitative traits subject to both drift and selection by considering random walks with some pull toward a particular state (39; 40; 41). Under our model, when *s ≈* 0 or *e*_*i*_ is very small, then 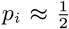, as in the neutral model. As *s* and *e*_*i*_ increase, *p*_*i*_ approaches 1, meaning that the direction of the effect is almost always in the positive direction.

### Analysis of pairwise comparisons in humans

#### Estimating *κ* from empirical population data

Estimating *κ* for a given pair of individuals using Eq. 1 requires (i) effect size differences for known loci to compute Δ, and (ii) the variance of the sum of the unknown effects, *σ*^2^. The genotype effect sizes can be ascertained from summary statistics of large genotype-phenotype datasets (see next section), from which we can compute the effect size differences (e.g., the added effect of one allele to the phenotype), denoted as *e*_*i*_. Next, we introduce a new parameter, 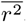, which denotes the overall contribution of known effects to the variance of phenotypic differences between pairs. 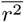 is similar to the proportion of explained phenotypic variance of PGS (usually denoted as *r*^2^), but refers only to those loci that are divergent between the two compared individuals; it is, therefore, expected to have similar values to *r*^2^ estimates computed using other means.

We assume that the measured differences in phenotypic values have been normalized and transformed to z-scores (i.e. the variance of the scaled phenotypic differences is one). To scale the units of the effect sizes to these standardized units, we define 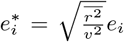, where *v*^2^ is the variance of in the PGS differences in the population. Thus, the effect sizes are scaled so that their overall contribution in units of the normalized phenotypic differences is 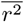. For a pair of individuals, we can now denote the overall predicted difference 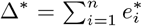, where *n* is the number of known effects that are divergent between the two individuals. To compute the variance of the sum of the unknown effects, we notice that the variance of the true phenotypic difference is composed of the sum of the variance explained by the known effects, 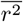, and the variance of unknown effects *σ*^2^; therefore, in the standardized units,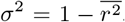. Using these standardized units, we can reformulate Eq. 1:

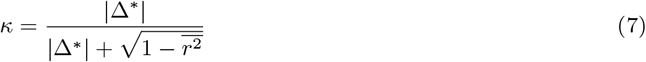

To apply this formulation to empirical data, we must estimate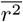. Below, we explore the option of estimating 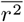 from the data by considering the fit to the theoretical expected relationship of *κ* and *P* (Eq. 4). We also computed, for comparison, the proportion of explained variance (*r*^2^) using the regression of the phenotypes on the PGSs.

#### Analysis of the UK Biobank

To test our approach on empirical data, we used the UK Biobank (UKB), a large dataset containing almost 500,000 genotyped individuals with associated phenotype data (12). We generated subsets of comparisons that have different levels of divergence: (i) sibling pairs with Northwestern European ancestry (within-family), (ii) pairs of individuals with Northwestern European ancestry (within-population), and (iii) pairs of individuals where each belongs to a different ancestry group, among European, East Asian, and African. Northwestern European ancestry was determined using the UKB Data-Field 22006. Our non-European groups were defined by demarcating clusters of genetically similar individuals that are distant from the European group on the PC1 and PC2 of the UKB PCA results from UKB Data-Field 22009 (Fig. S6). The two clusters were labeled as East Asian and African based on the majority of self-identifications of individuals from these groups as reported in UKB Data-Field 21000. These groups included 1,794 and 3,091 individuals, respectively.

To compute *κ* values, we first generated GWAS results for a number of continuous traits: body-mass index (UKB Data-Field 21001); systolic blood pressure (UKB Data-Field 4080); heel bone mineral density (UKB Data-Field 3148); standing height, referred to as “height” (UKB Data-Field 50); hip circumference (UKB Data-Field 49); and basal metabolic rate, referred to as “metabolic rate” (UKB Data-Field 23105). We included variants with high-quality imputation scores (imputation INFO scores *≥* 0.8) from the UKB imputed genotype release version 3 (12); this yielded roughly 30 million variants. The discovery dataset included individuals with Northwestern European ancestry, excluding 20,000 (10,000 female, 10,000 male) individuals as a validation subset. We generated single-variant association results using SAIGE v1.1.6.3 (42). We used 280,628 markers to fit the null linear mixed model, and age, sex, and the first ten genetic PCs as covariates. To generate PGSs, GWAS results were filtered with a fixed P-value threshold of P-value *≤* 0.01 and minor allele count threshold of MAC *≥* 20. We used PRSice-2 to compute PGSs for all individuals (43). *κ* values were computed for all same-sex pairs from our validation subset as detailed above in *Estimating κ from empirical data*. For each pair, we compared the sign of the PGS difference and the true direction of phenotype difference as reported in the UKB.

To compare our results to the theoretical expectation of the relationship between *κ* and prediction accuracy, we binned comparisons according to their *κ* values, and computed the proportion of correct predictions in each bin. To estimate 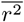 from the data, we computed *κ* values for a range of 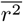 values, and selected the value of 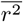 that yielded the least sum of absolute distances between the proportion of correct predictions for each bin and the theoretical expectation, weighted by the number of comparisons per bin (Table S2).

PGSs are known to have poor transferability between genetically distinct populations. To test the effect of PGS transferability on our model fit, we used the PGSs from the European ancestry group in Yengo *et al*. (15) to evaluate our predictions in non-European pair comparisons, using the ancestry subsets indicated above (1,794 individuals with East Asian ancestry for EAS–EAS comparisons, and 3,091 individuals with African ancestry for AFR–AFR comparisons), relative to our pairwise predictions in the European group with the same PGSs (20,000 individuals for EUR–EUR comparisons). Note that in the EUR–EUR comparisons the Yengo *et al*. (15) PGSs included the tested individuals, but these individuals constitute a very small portion of the overall European population analyzed in this study.

We also generated predictions for a number of common diseases reported in the UKBB according to the following ICD10 codes: asthma (J45), type 2 diabetes (E11), hypertension (I10) and hypothyroidism (E03). ICD10 codes were retrieved from UKB Data-Field 41270 (diagnoses). For each disease, we generated single-variant association results using SAIGE2 (42) for binary traits with default parameters. The discovery dataset included individuals with Northwestern European ancestry (as defined by UKB Datafield 22006), excluding 10,000 samples, 5,000 controls and 5,000 cases, as a validation subset. PGSs were generated and *κ* estimated as for the continuous traits. For each case-control pair, correct prediction was recorded whenever the PGS for the disease risk was higher in the case individual.

#### Analysis of population and species datasets

To evaluate our approach in cases where the compared genomes are highly diverged, we examined datasets from several species. In the stickleback QTL mapping dataset (17), we compared a marine population (treated in our analysis as the *phenotyped* population) and four freshwater populations (treated as *unphenotyped*). The compared populations likely diverged less than 12,000 years ago (17). We investigated 27 morphological phenotypes (measurements of shape landmark coordinates), resulting in four pairwise comparisons of 27 phenotypes. Because not all phenotypes had significant QTLs in each population, some of the comparisons (three out of four populations) included fewer than 27 predictions (Fig. 4c). Here, because the raw data was not available, we could not exclude the compared individuals when computing effect sizes; however, because these loci are largely fixed between the populations, this is not expected to affect the results.

In the mouse QTL mapping dataset (19), we compared a wild-derived inbred laboratory house mouse strain and the Gough island house mouse subpopulation. These populations diverged in the 19^th^ century. Two phenotypes (weight and growth rate) were measured across 16 weeks, resulting in a pairwise comparison of 2 *×* 16 phenotypes. We then computed average prediction accuracy across the 16 time points for each of the two phenotypes.

In the daisy QTL mapping dataset (18), we compared two daisy species (*Senecio aethnensis*, and *Senecio chrysanthemifolius*) that have likely diverged within the last 176,000 years (Brennan et al., 2016). For one phenotype out of 13, a prediction could not be made because the sum of the known effects was 0.

The Neanderthal and chimpanzee datasets (6) included comparisons of DNA methylation maps between modern humans (treated as the *phenotyped* population) and Neanderthals and chimpanzees. Because these analyses do not contain effect sizes, they were limited to phenotypes for which the loci with the largest differences in methylation levels showed unidirectionality (likely resulting in high Δ values, and therefore high *κ* values). These analyses predicted the phenotypic direction for 33 Neanderthal phenotypes and 22 chimpanzee phenotypes. We list here the prediction accuracy as reported in (6).

## Acknowledgments

We thank David Reich for the original idea to test this approach with a model, and Dmitri Petrov, Hunter Fraser, Noah Rosenberg, Arbel Harpak, Guy Sella, Yuval Simons, Liran Carmel, Jaehee Kim, John (Tony) Capra, Moi Exposito-Alonso, and members of the Fraser, Petrov, Rosenberg, Greenbaum, and Gokhman labs for input. This research was partially supported by the Israeli Council for Higher Education (CHE) via the Weizmann Data Science Research Center, a research grant from the Center for New Scientists at the Weizmann Institute of Science, and the Kahn Family Research Center for Systems Biology of the Human Cell. SC was supported by the National Institutes of Health (grant R01HG011711). This research has been conducted using data from UK Biobank, a major biomedical database, UK Biobank project ID 26664.

## References

[1] Rosenberg, N. A., Edge, M. D., Pritchard, J. K. & Feldman, M. W. Interpreting polygenic scores, polygenic adaptation, and human phenotypic differences. Evolution, Medicine, and Public Health 2019, 26–34 (2019).

[2] Young, A. I., Benonisdottir, S., Przeworski, M. & Kong, A. Deconstructing the sources of genotype-phenotype associations in humans. Science 365, 1396–1400 (2019).

[3] Dittmar, E. L., Oakley, C. G., Conner, J. K., Gould, B. A. & Schemske, D. W. Factors influencing the effect size distribution of adaptive substitutions. Proceedings of the Royal Society B: Biological Sciences 283, 20153065 (2016).

[4] Orr, H. A. The genetic theory of adaptation: a brief history. Nature Reviews Genetics 6, 119–127 (2005).

[5] Scheben, A. & Edwards, D. Towards a more predictable plant breeding pipeline with CRISPR/Cas-induced allelic series to optimize quantitative and qualitative traits. Current Opinion in Plant Biology 45, 218–225 (2018).

[6] Gokhman, D. et al. Reconstructing Denisovan Anatomy Using DNA Methylation Maps. Cell 179, 180–192.e10 (2019).

[7] Thudi, M. et al. Genomic resources in plant breeding for sustainable agriculture. Journal of Plant Physiology 257, 153351 (2021).

[8] Karavani, E. et al. Screening Human Embryos for Polygenic Traits Has Limited Utility. Cell 179, 1424–1435.e8 (2019).

[9] Lello, L., Raben, T. G. & Hsu, S. D. Sibling validation of polygenic risk scores and complex trait prediction. Scientific Reports 10, 13190 (2020).

[10] Widen, E., Lello, L., Raben, T. G., Tellier, L. C. & Hsu, S. D. Polygenic Health Index, General Health, and Pleiotropy: Sibling Analysis and Disease Risk Reduction. Scientific Reports 12, 18173 (2022).

[11] Palmer, D. S. et al. Analysis of genetic dominance in the UK Biobank. Science 379, 1341–1349 (2023).

[12] Bycroft, C. et al. The uk biobank resource with deep phenotyping and genomic data. Nature 562, 203–209 (2018).

[13] Berg, J. J. et al. Reduced signal for polygenic adaptation of height in UK biobank. eLife 8, e39725 (2019).

[14] Martin, A. R. et al. Clinical use of current polygenic risk scores may exacerbate health disparities. Nature Genetics 51, 584–591 (2019).

[15] Yengo, L. et al. A saturated map of common genetic variants associated with human height. Nature 610, 704–712 (2022).

[16] Hayward, L. K. & Sella, G. Polygenic adaptation after a sudden change in environment. eLife 11, e66697 (2022).

[17] Rogers, S. M. et al. Genetic signature of adaptive peak shift in threespine stickleback. Evolution 66, 2439–2450 (2012).

[18] Brennan, A. C., Hiscock, S. J. & Abbott, R. J. Genomic architecture of phenotypic divergence between two hybridizing plant species along an elevational gradient. AoB Plants 8, plw022 (2016).

[19] Gray, M. M. et al. Genetics of rapid and extreme size evolution in Island mice. Genetics 201, 213–228 (2015).

[20] Gokhman, D. et al. Human–chimpanzee fused cells reveal cis-regulatory divergence underlying skeletal evolution. Nature Genetics 53, 467–476 (2021).

[21] Hill, W. G., Goddard, M. E. & Visscher, P. M. Data and theory point to mainly additive genetic variance for complex traits. PLoS Genetics 4, e1000008 (2008).

[22] Hivert, V. et al. Estimation of non-additive genetic variance in human complex traits from a large sample of unrelated individuals. bioRxiv 2020.11.09.375501 (2020).

[23] Pazokitoroudi, A., Chiu, A. M., Burch, K. S., Pasaniuc, B. & Sankararaman, S. Quantifying the contribution of dominance effects to complex trait variation in biobank-scale data. bioRxiv 2020.11.10.376897 (2020).

[24] Exposito-Alonso, M., Wilton, P. & Nielsen, R. Non-additive polygenic models improve predictions of fitness traits in three eukaryote model species. bioRxiv 2020.07.14.194407 (2020).

[25] Mackay, T. F. Epistasis and quantitative traits: Using model organisms to study gene-gene interactions. Nature Reviews Genetics 15, 22–33 (2014).

[26] Harpak, A. & Przeworski, M. The evolution of group differences in changing environments. PLoS Biology 19, e3001072 (2021).

[27] Yair, S. & Coop, G. Population differentiation of polygenic score predictions under stabilizing selection. Philosophical Transactions of the Royal Society B: Biological Sciences 377 (2022).

[28] Polyakov, A. et al. Polygenic risk score for embryo selection-not ready for prime time. Human Reproduction 37, 2229–2236 (2022).

[29] Treff, N. R., Savulescu, J., de Melo-Martín, I., Shulman, L. P. & Feinberg, E. C. Should preimplantation genetic testing for polygenic disease be offered to all – or none? Fertility and Sterility 117, 1162–1167 (2022).

[30] Lencz, T. et al. Concerns about the use of polygenic embryo screening for psychiatric and cognitive traits 9, 838–844 (2022).

[31] Siermann, M. et al. Limitations, concerns and potential: attitudes of healthcare professionals toward preimplantation genetic testing using polygenic risk scores. European Journal of Human Genetics 31, 1133–1138 (2023).

[32] Forzano, F. et al. The use of polygenic risk scores in pre-implantation genetic testing: an unproven, unethical practice. European Journal of Human Genetics 30, 493–495 (2022).

[33] Treff, N. R. et al. Preimplantation genetic testing for polygenic disease relative risk reduction: Evaluation of genomic index performance in 11,883 adult sibling pairs. Genes 11, 648 (2020).

[34] Lencz, T. et al. Utility of polygenic embryo screening for disease depends on the selection strategy. eLife 10, e64716 (2021).

[35] Dempster, E. R. & Lerner, I. M. Heritability of threshold characters. Genetics 35, 212–236 (1950).

[36] Turley, P. et al. Problems withusing polygenic scores to select embryos. The New England Journal of Medicine 385, 78–86 (2021).

[37] Simons, Y. B., Mostafavi, H., Smith, C. J., Pritchard, J. K. & Sella, G. Simple scaling laws control the genetic architectures of human complex traits. bioRxiv 2022.10.04.509926 (2022).

[38] Orr, H. A. The population genetics of adaptation: The distribution of factors fixed during adaptive evolution. Evolution 52, 935–949 (1998).

[39] Bedford, T. & Hartl, D. L. Optimization of gene expression by natural selection. Proceedings of the National Academy of Sciences 106, 1133–1138 (2009).

[40] Butler, M. A. & King, A. A. Phylogenetic comparative analysis: a modeling approach for adaptive evolution. The American Naturalist 164, 683–695 (2004).

[41] Hansen, T. F. Stabilizing selection and the comparative analysis of adaptation. Evolution 51, 1341–1351 (1997).

[42] Zhou, W. et al. Efficiently controlling for case-control imbalance and sample relatedness in large-scale genetic association studies. Nature Genetics 50, 1335–1341 (2018).

[43] Choi, S. W. & O’Reilly, P. F. Prsice-2: Polygenic risk score software for biobank-scale data. Gigascience 8, giz082 (2019).

