## Supporting Information for "Predicting the direction of phenotypic difference"

#### Independence of the distribution of $\kappa$ from the effect size distribution

Here, we derive the probability distribution function of $\kappa$ to show that it is independent of the effect size distribution. We denote the number of known effects as $n$ and unknown effects as $m$, and therefore the fraction of known effects is $r = \frac{n}{n+m}$. The known effects sizes are denoted as $e_1, \cdots, e_n$ and the unknown effects as effects as $e_{n+1}, \ldots, e_{n+m}$. We assume that the effects are independent, identically-distributed random variables with $E[e_i] = 0$ and $Var[e_i] = \sigma_E^2$.

As in the main text, we denote the sum of the known effects as $\Delta' = \sum_{i=1}^{n} e_i$. Because the effects are identical and independent, we can use the central limit theorem (assuming $n$ is large) to conclude that $\Delta'$ is normally distributed with zero mean and variance $Var[\Delta'] = n\sigma_E^2$. We take the positive sum $\Delta = |\Delta'|$, which is half-normal, meaning that it has a probability density that is twice that of the normal distribution:

$$f(\Delta) = \sqrt{\frac{2}{n\pi}} \cdot \frac{1}{\sigma_E} \cdot e^{-\frac{\Delta^2}{2n\sigma_E^2}} \tag{S1}$$

Similarly, the sum of the unknown effects is denoted as $\Sigma = \sum_{i=n+1}^{n+m} e_i$, with variance which we denote as $\sigma^2 = m\sigma_E^2$.

We can now derive the PDF of our statistic $\kappa = \frac{\Delta}{\Delta+\sigma}$ using transformation of random variables ($\kappa(\Delta)$ is strictly increasing and differentiable):

$$f(\kappa) = f(\Delta)\frac{d\Delta}{d\kappa}. \tag{S2}$$

We can rearrange the formulation of $\kappa$ to obtain $\Delta = \frac{\kappa\sigma}{1-\kappa}$ and find the derivative,

$$\frac{d\Delta}{d\kappa} = \frac{\sigma}{(1-\kappa)^2}. \tag{S3}$$

Substituting Eq. S1 and Eq. S3 into Eq. S2, we obtain:

$$f(\kappa) = \sqrt{\frac{2}{n\pi}} \cdot \frac{1}{\sigma_E} \cdot e^{-\frac{\Delta^2}{2n\sigma_E^2}} \cdot \frac{\sigma}{(1-\kappa)^2}. \tag{S4}$$

We can now substitute $\sigma^2 = m\sigma_E^2$ and $\Delta = \frac{\kappa\sigma}{1-\kappa}$ to obtain the PDF of $\kappa$ explicitly:

$$f(\kappa) = \sqrt{\frac{2}{n\pi}} \cdot \frac{1}{\sigma_E} \cdot e^{-\frac{\left(\frac{\kappa\sqrt{m}\sigma_E}{1-\kappa}\right)^2}{2n\sigma_E^2}} \cdot \frac{\sqrt{m}\sigma_E}{(1-\kappa)^2} = \sqrt{\frac{2}{\pi}} \cdot \sqrt{\frac{m}{n}} \cdot \frac{1}{(1-\kappa)^2} e^{-\frac{m}{2n}(\frac{\kappa}{1-\kappa})^2}. \quad \text{(S5)}$$

We can reformulate the PDF using the fraction of known effects $r$, noting that $\frac{m}{n} = \frac{1-r}{r}$:

$$f(k) = \sqrt{\frac{2}{\pi}} \cdot \sqrt{\frac{1-r}{r}} \cdot \frac{1}{(1-\kappa)^2} e^{-\frac{1-r}{2r}(\frac{\kappa}{1-\kappa})^2}. \quad \text{(S6)}$$

Notice that the $\sigma_E$ terms cancels out in Eq.S5, implying that the PDF of $\kappa$ is independent of the effect size distribution. To demonstrate this point, and to investigate a scenario where $n$ and $m$ are not infinite as required for the use of the central limit theorem above, we simulated two different effect size distribution ($r = 0.5$, $n + m = 10,000$) with different paramaterizations to show that the $\kappa$ distributions are independent of the effect size distribution (Fig. S1).

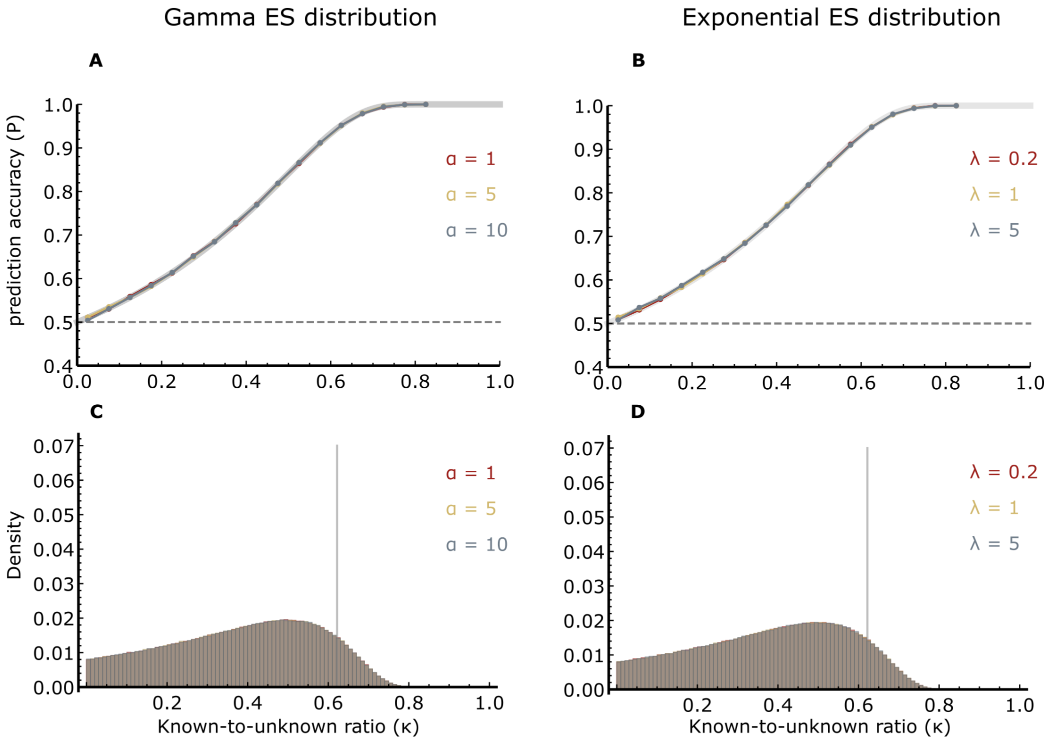


**Figure S1:** Simulation results with effect sizes sampled from different distributions. (A) The relationship between $\kappa$ and prediction accuracy ($P$) when the effect sizes are sampled from a Gamma distribution with a scale parameter $\beta = 1$, and three different shape parameters, $\alpha = 1$, 5, and 10. (B) The relationship between $\kappa$ and $P$ when the effect sizes are sampled from an Exponential distribution with three different rate parameters, $\lambda = 0.2$, 1 and 5. (C) The distribution of $\kappa$ values generated by the simulation with Gamma effect size distribution described in panel A. (D) The distribution of $\kappa$ values generated by the simulation with Exponential effect size distribution described in panel B. In the top panels, the gray curve denotes the theoretical expectation of the relationship between $\kappa$ and $P$. In the bottom panels, the vertical grey line denotes the $\kappa$ values that corresponds to $P = 0.95$ based on the theoretical expectation. In all simulations, 10,000 effect were simulated, where it is assumed that 50% of them are known. For each distribution parameter, $10^6$ repeats were simulated. Note that the results with different effect size distributions are very similar (the curves in the top panels and distributions in the bottom panels are on top of each other).

### Simulation results

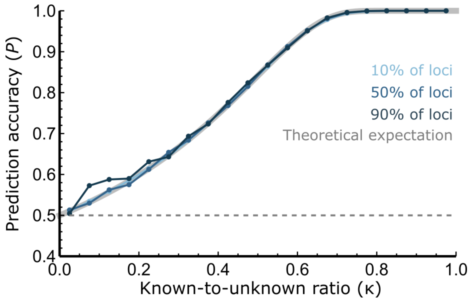


**Figure S2:** Simulation results of the relationship between $\kappa$ and $P$ for the case when the known effect sizes are the largest ones. We simulated three scenarios, with different portions of known effects, 10%, 50% and 90%. In all simulations, 10,000 effect were simulated. For each portion of known effects, $10^6$ repeats were simulated.

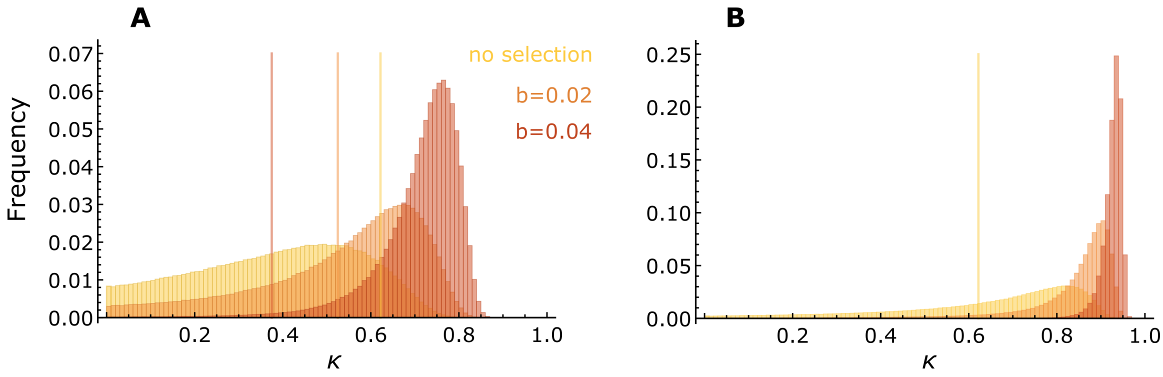


**Figure S3:** Distribution of $\kappa$ values generated by simulations with selection. We simulated three scenarios, with different strength of selection parameters, $s = 0$, 0.02 and 0.04. (A) Known effects are assumed to be a random sample of all effect sizes. (B) known effects are assumed to be the largest effects. The vertical lines denote the $\kappa$ values required for prediction accuracy of $P > 0.95$ . In all simulations, 10,000 effects were randomly drawn from a standard normal distribution, and 50% of the effects were assumed to be known. For each strength of selection, $10^6$ repeats were simulated.

### Analysis of human data (UK Biobank)

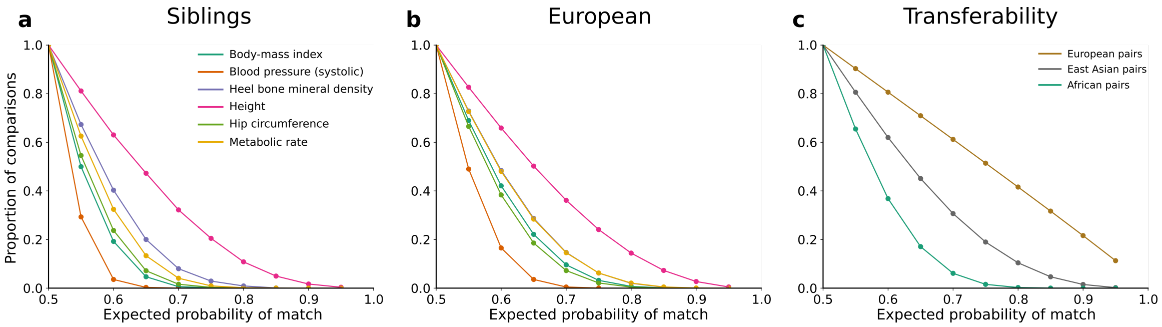


**Figure S4:** Proportion of pairwise comparisons with prediction success greater than a given value, for the analyses presented in Figure 3 of the main text. For each pairwise comparison, its $\kappa$ value was translated to an expected prediction accuracy value using the theoretical expectation (Eq. 2). (a)–(c) The curves show the cumulative expected prediction accuracy, computed using all pairwise comparisons used in the analyses of Figure 3. The colors in panel b correspond to the legend in panel a.

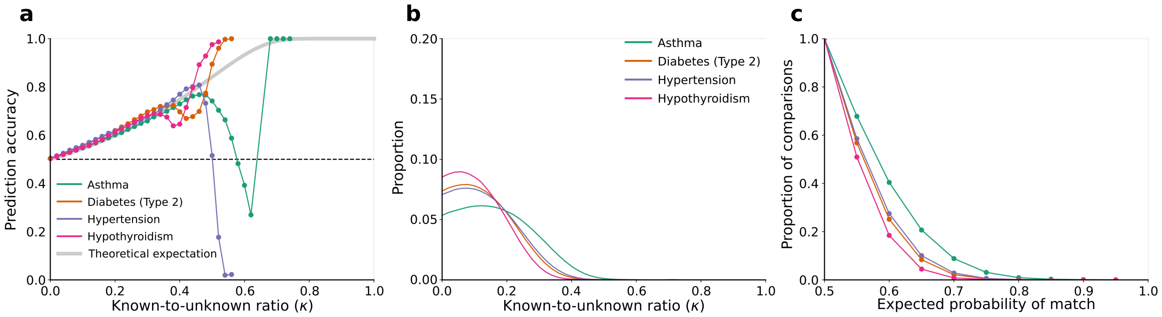


**Figure S5:** Analysis of disease phenotypes in individuals with Northwest European ancestry in the UK Biobank. (a) Relationship between the known-to-unknown ratio ($\kappa$) and prediction accuracy ($P$). (b) The distribution of $\kappa$ values for all pairwise comparisons. (c) Proportion of pairwise comparisons with prediction success greater than a given value.

| Trait | EUR $r^2$ (F/M) | EUR $\overline{r^2}$ | Siblings $r^2$ (F/M) | Siblings $\overline{r^2}$ |
|---|---|---|---|---|
| Body-mass index | 0.07 / 0.09 | 0.09 | 0.07 / 0.07 | 0.04 |
| Blood pressure (systolic) | 0.02 / 0.03 | 0.03 | 0.03 / 0.01 | 0.01 |
| Heel bone mineral density | 0.10 / 0.06 | 0.12 | 0.10 / 0.08 | 0.08 |
| Hip circumference | 0.06 / 0.07 | 0.08 | 0.05 / 0.05 | 0.04 |
| Metabolic rate | 0.08 / 0.11 | 0.12 | 0.09 / 0.10 | 0.06 |
| Standing height | 0.22 / 0.20 | 0.25 | 0.21 / 0.21 | 0.22 |

**Table S1:** Comparison of explained variance ($r^2$) of the regression of PGS values and phenotypes (computed separately for same-sex subsets) and inferred $\overline{r^2}$ values, in the European ancestry (EUR) and siblings subsets.

| Analysis | Trait/Comparison | $\overline{r^2}$ |
|---|---|---|
| Siblings traits | Body-mass index | 0.04 |
| Siblings traits | Systolic blood pressure | 0.01 |
| Siblings traits | Heel bone mineral density | 0.08 |
| Siblings traits | Hip circumference | 0.04 |
| Siblings traits | Metabolic rate | 0.06 |
| Siblings traits | Standing height | 0.22 |
| EUR-EUR traits | Body-mass index | 0.09 |
| EUR-EUR traits | Systolic blood pressure | 0.03 |
| EUR-EUR traits | Heel bone mineral density | 0.12 |
| EUR-EUR traits | Hip circumference | 0.08 |
| EUR-EUR traits | Metabolic rate | 0.12 |
| EUR-EUR traits | Standing height | 0.25 |
| Transferability | EUR–EUR (Height) | 0.52 |
| Transferability | EAS–EAS (Height) | 0.21 |
| Transferability | AFR–AFR (Height) | 0.07 |
| EUR-EUR diseases | Asthma | 0.04 |
| EUR-EUR diseases | Diabetes (Type 2) | 0.05 |
| EUR-EUR diseases | Hypertension | 0.05 |
| EUR-EUR diseases | Hypothyroidism | 0.09 |

**Table S2:** $\overline{r^2}$ values inferred in the analyses in Figure 3 in the main text and Figure S5.

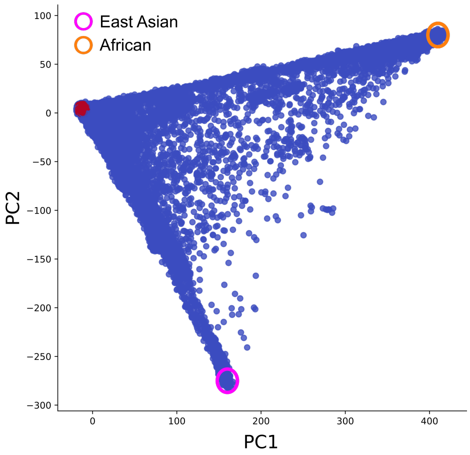


**Figure S6:** PCA of the UK Biobank individuals used for definition of populations in the analysis presented in Figure 3c and 3f in the main text. Individuals self-identified as ‘White British’ and clustered with Northwest European ancestry (according to UKB Data-Field 22006) are marked in red. For between-population comparisons, we defined two groups in the PCA that are distant from this European group, one with $1,794$ genetically similar individuals, the majority of whom self-identified with ethnicity associated with the geographic region of East Asia (samples encircled in pink), and one with $3,091$ genetically similar individuals, the majority of whom self-identified with ethnicity associated with the geographic region of Africa (samples encircled in orange).

### Derivation of prediction accuracy for specific cases

#### Siblings

We can use this formulation to also derive the prediction accuracy $P$ for the case when the two compared individuals are siblings. In this case, we need to consider the shared genetic component and the shared environmental component. We therefore model the true phenotypic values of the two compared individuals as $y_1 = p_1 + \overline{g_1} + \overline{e_1}$ and $y_2 = p_2 + \overline{g_2} + \overline{e_2}$, where the PGSs are $p_i \sim N(0, r^2)$ as before, $\overline{g_i}$ is the genetic component that is not modeled in the score and is distributed as $\overline{g_i} \sim N(h^2 - r^2)$ with $h^2$ being the overall narrow-sense heritability of the phenotype in the population, and $\overline{e_i}$ is the non-genetic component which is distributed $e_i \sim N(0, 1 - h^2)$. We then consider the shared genetic and environmental components by defining $\overline{g_i} = g_i + g_s$ and $\overline{e_i} = e_i + e_s$, where $g_s$ and $e_s$ are the shared genetic and environmental components, respectively, and $g_i$ and $e_i$ are unique to each sibling. In siblings, the correlation of the genetic components is 0.5, and therefore $g_i \sim N(0, \frac{h^2 - r^2}{2})$. For the environment, we define the shared environmental variance as $c^2$, and therefore $e_i \sim N(0, 1 - h^2 - c^2)$ (1). We can now derive the prediction accuracy when comparing the two siblings given their PGS difference $d$:

$$P = Prob(y_1 > y_2) = Prob(p_1 + g_1 + g_s + e_1 + e_s > p_2 + g_2 + g_s + e_2 + e_s) = Prob\Big(g_2 - g_1 + e_2 - e_1 < d\Big). \tag{S7}$$

Assuming that the $g_i$'s and $e_i$'s are all pairwise independent, the variance of $(g_2 - g_1) + (e_2 - e_1)$ is $(h^2 - r^2) + 2(1 - h^2 - c^2) = 2 - h^2 - r^2 - 2c^2$. Therefore, the prediction accuracy of sibling comparisons can is given by

$$P = \Phi\Big(\frac{d}{\sqrt{2 - h^2 - r^2 - 2c^2}}\Big). \tag{S8}$$

Note that, given that $r^2 < h^2$, as the PGS cannot explain more variance than the heritability, we have $2 - h^2 - r^2 - 2c^2 < 2 - r^2 - r^2 - 2c^2 < 2 - 2r^2$. Thus, $\Phi\Big(\frac{d}{\sqrt{2-h^2-r^2-2c^2}}\Big) > \Phi\Big(\frac{d}{\sqrt{2(1-r^2)}}\Big)$. This is expected, because a given PGS difference is more informative on phenotypic differences between siblings compared to unrelated pairs.

#### Diseases

For diseases, we can apply the widely-used liability threshold model (2). Under the model, $y$ is the overall liability to develop the disease. As for continuous traits, $y = p + e$, $y \sim N(0, 1)$, $p \sim N(0, r^2)$, and $e \sim N(0, 1 - r^2)$ represents both unmodeled genetic factors and non-genetic factors contributing to the liability. An individual is affected whenever $y > T$, where $T$ is the liability threshold, equal to $T = \Phi^{-1}(1-K)$, where $K$ is the disease prevalence and $\Phi^{-1}$ is the inverse normal CDF.

Consider next two (unrelated) individuals, having PGSs $p_1$ and $p_2$, with $p_1 - p_2 = d$. We assume that $p_1, p_2, e_1, e_2$ are all pairwise independent. In our simulations, we only considered discordant pairs of individuals, where one is affected and the other is unaffected. Given $d > 0$, the prediction accuracy can be written as the conditional probability

$$P = \frac{Prob(\text{1 affected; 2 unaffected})}{Prob(\text{1 affected; 2 unaffected}) + Prob(\text{1 unaffected; 2 affected})} \tag{S9}$$

We show below how to compute $Prob$(1 affected; 2 unaffected). The calculation of the second term is analogous. We first condition on the PGSs $p_1$ and $p_2$.

$$\begin{aligned} Prob(\text{1 affected; 2 unaffected}|p_1,p_2) &= Prob(y_1 > T, y_2 < T|p_1,p_2) \\ &= Prob(p_1 + e_1 > T, p_2 + e_2 < T) \\ &= Prob(e_1 > T - p_1, e_2 < T - p_2) \\ &= \left[1 - \Phi\left(\frac{T-p_1}{\sqrt{1-r^2}}\right)\right] \Phi\left(\frac{T-p_2}{\sqrt{1-r^2}}\right). \end{aligned} \tag{S10}$$

Next, the individuals are randomly selected from the population (such that $p_1$ and $p_2$ are normal with zero mean and variance $r^2$). Integrating over all $p_1$ and $p_2$, taking into account that the difference between them is $d$, we obtain

$$\begin{aligned} &= Prob(\text{1 affected; 2 unaffected}) \\ &= \int_{-\infty}^{\infty}\int_{-\infty}^{\infty} \frac{1}{r}\phi\left(\frac{p_1}{r}\right)\frac{1}{r}\phi\left(\frac{p_2}{r}\right)\delta(p_1-p_2-d)\left[1-\Phi\left(\frac{T-p_1}{\sqrt{1-r^2}}\right)\right]\Phi\left(\frac{T-p_2}{\sqrt{1-r^2}}\right)dp_2dp_1 \\ &= \int_{-\infty}^{\infty} \frac{1}{r}\phi\left(\frac{p_1}{r}\right)\frac{1}{r}\phi\left(\frac{p_1-d}{r}\right)\left[1-\Phi\left(\frac{T-p_1}{\sqrt{1-r^2}}\right)\right]\Phi\left(\frac{T-p_1+d}{\sqrt{1-r^2}}\right)dp_1 \\ &= \frac{1}{r}\int_{-\infty}^{\infty} \phi(t)\phi\left(t-d/r\right)\left[1-\Phi\left(\frac{T-rt}{\sqrt{1-r^2}}\right)\right]\Phi\left(\frac{T-rt+d}{\sqrt{1-r^2}}\right)dt, \end{aligned} \tag{S11}$$

where $\phi$ is the normal PDF and we changed variables $t = p_1/r$. In practice, the resulting expression has very similar shape to a (non-standard) normal CDF.

### References

[1] Lakhani, C. M. *et al.* Repurposing large health insurance claims data to estimate genetic and environmental contributions in 560 phenotypes. *Nature Genetics* **51**, 327–334 (2019).

[2] Visscher, P. M. & Wray, N. R. Concepts and Misconceptions about the Polygenic Additive Model Applied to Disease. *Human Heredity* **80**, 165–170 (2016).
